# *In vitro* replicative potential of an HIV-1/MO intergroup recombinant virus compared to HIV-1/M and HIV-1/O parental viruses

**DOI:** 10.1101/2022.10.24.513636

**Authors:** Alice Moisan, Fabienne De Oliveira, Manon Mabillotte, Elodie Alessandri-Gradt, Thomas Mourez, Jean-Christophe Plantier

**Affiliations:** Normandie Univ, UNIROUEN, UNICAEN, Inserm UMR 1311 DYNAMICURE, CHU Rouen, Department of Virology, Rouen, France; Normandie Univ, UNIROUEN, UNICAEN, Inserm UMR 1311 DYNAMICURE, 76000 Rouen, France

**Author notes:** Alternative corresponding author.

**Keywords:** Replicative potential, HIV-1/MO recombinant, Cloning, Recombination, fitness, URF_MO

## Abstract

HIV-1 is characterized by high genetic diversity and genetic recombination is one of the major evolution processes. Despite their great genetic divergence, HIV-1 group M, pandemic, and group O, endemic in Cameroon, can generate HIV-1/MO intergroup recombinants. The current description of 19 HIV-1/MO recombinant forms (URF_MO) raises the question of a possible benefit of the recombination and the modalities of their emergence.

Therefore, the objectives of this work were to study *in vitro* the replicative potential of HIV-1/MO recombinant forms.

The replicative potential was analyzed, based on a simple recombination pattern, [O_gag/pol_-M_env_], harboring a breakpoint in Vpr, due to a recombination hotspot in this region. For this, a chimeric infectious molecular clone (IMC) was synthesized from HIV-1/M subtype B and HIV-1/O subgroup T and recombinant viruses were obtained by transfection and co-culture. To compare the replicative potential of recombinant viruses with HIV-1/M and HIV-1/O parental viruses, two markers were monitored in culture supernatants: Reverse Transcriptase (RT) activity and P24 antigen concentration. The results showed a superiority of the group M parental virus compared to group O for both markers. In contrast, for the HIV-1/OM recombinant virus, RT activity data did not overlap with the concentration of P24 antigen, suggesting a hybrid behavior of the recombinant, in terms of enzyme activity and P24 production.

These results highlighted many hypotheses about the impact of recombination on replicative potential and demonstrated once again the significant plasticity of HIV genomes and their infinite possibility of evolution.

**Importance:** HIV-1/M and HIV-1/O can generate HIV-1/MO intergroup recombinants. The current description of 19 URF_MO raises the question of a possible benefit of recombination in terms of emergence. The objectives of this work were to study *in vitro* the replicative potential of HIV-1/MO recombinant viruses. For this, a chimeric infectious molecular clone (IMC) generated from HIV-1/M subtype B and HIV-1/O subgroup T and recombinant viruses were obtained by transfection and co-culture. To compare the replicative potential of recombinant viruses with HIV-1/M and HIV-1/O parental viruses, RT activity and P24 antigen concentration were monitored in culture supernatants. A superiority of the group M parental virus compared to group O was observed for both markers whereas a hybrid behavior in terms of enzyme activity and P24 production was found for the recombinant virus. These results demonstrated once again the significant plasticity of HIV genomes and their infinite possibility of evolution.

## INTRODUCTION

HIV-1 is characterized by high genetic diversity, due to its simian origins and replication mode, enhanced by recombination events. Despite the great genetic divergence between HIV-1 group M (HIV-1/M), pandemic, and group O (HIV-1/O), endemic in Cameroon, HIV-1/M+O dual infections can generate HIV-1/MO inter-group recombinants [1]. Over the past 20 years, several case reports and epidemio-molecular studies have shown the ability of HIV-1/M and /O to superinfect a patient already infected with a viral form from another group (1 case) [2]; to replicate in the context of double infections (17 cases) [3–8]; and to generate intergroup recombinant forms capable of replicating in the absence (10 cases) [5, 7, 13, 14] or in the presence of one or both parental forms (9 cases) [5, 7–12]. Phylogenetic analysis showed that the 19 recombinant forms corresponded to 19 unique HIV-1/MO recombinant forms (URF_MO). The analysis of the genomic profiles and of the breakpoint frequency revealed hotspots in the “central” accessory genes (*vif*, *vpr*, *vpu*), Long Terminal repeat (LTR) regions, reverse transcriptase (RT) and gp41, and no recombination event in protease, gp120 and nef [15]. This work also highlighted a variable degree of complexity in profiles, with on average two to three breakpoints per recombinant genome [15]. The current description of these 19 URF_MOs, raises the question of a possible benefit of recombination and the modalities of their emergence. The evolution of the genetic diversity of HIV-1/M, of which the number of recombinant forms, both circulating recombinant forms (CRF) and URFs, has escalated since the beginning of the epidemic, clearly shows the interest of studying URF_MOs. Indeed, studies have suggested a better adaptation and a better fitness of intragroup M recombinants compared to parental forms and have highlighted preferential recombination profiles [16–18]. Therefore, data collected from the URF_MOs currently described *in vivo* and the presence of HIV-1/MO recombinant forms in patients with neither of the two original parental forms suggest the possible existence of phenotypic properties conferring a selective advantage of these HIV-1/MO recombinant forms, compared to HIV-1/M and HIV-1/O parental forms. Moreover, this emergence of multiple URF_MOs could lead over time to the successful emergence of circulating forms (i.e. a CRF_MO).

Regarding the relative replicative fitness, a pairwise growth competition experiment showed that HIV-1 group O was 10 to 100-fold-less fit than group M forms, when competed in peripheral blood mononuclear cells (PBMC), and depending on the diversity of the forms studied. These results suggested a lesser replicative fitness of group O, contributing to the low prevalence and limited geographical spread of HIV-1/O in the human population although it emerged several years ago [19, 20]. The phenomenon of M/O inter-group recombination could thus rebalance this “pyramid of fitness” and confer a replicative advantage to an HIV-1/MO recombinant form compared to one, the other or both of the parental forms. To date, only one study has investigated the replicative potential of an HIV-1/MO recombinant form and the results indicated a better *in vitro* adaptive capacity (fitness) of the recombinant virus compared to the HIV-1/M parental virus, suggesting a potential better spread and therefore a selective advantage conferred by recombination, in the absence of selection pressure [12].

The aim of this work was to investigate *in vitro* the viral properties conferred by recombination, on the replicative potential of an HIV-1/MO recombinant virus compared to both of the HIV-1/M and /O parental viruses, from which it was derived.

## MATERIALS AND METHODS

### Parental infectious molecular clones and chimeric infectious molecular clones

#### Selection of parental forms and positioning of breakpoint

Given the reverse genetics approach that we aimed to implement, two parental infectious molecular clones (PIMC) were used as parental viruses: pRBF206 and p89.6. RBF206 is an HIV-1/O subgroup T form isolated and characterized in our laboratory, with an R5 tropism. The corresponding PIMC pRBF206 was obtained as previously described [26]. For HIV-1/M, 89.6 belongs to the subtype B and shows an X4 tropism. The PIMC p89.6 was obtained from the National Institute for Biological Standards and Control (NIBSC) Centralized Facility for AIDS Reagent (CFAR). To generate the chimeric infectious molecular clones (CIMC), we first designed the recombination patterns and chose a breakpoint, on the basis of our previous study [15]. The recombination patterns were chosen to reproduce the two major chimeric genome profiles described *in vivo*: [O_gag/pol_-M_env_] and [M_gag/pol_-O_env_], and to generate the recombinant viruses HIV-1/OM and HIV-1/MO, respectively. Given the recombination hotspot identified *in vivo* [15], the breakpoint was positioned in a conserved region identified in the *vpr* gene of HIV-1/M and HIV-1/O genomes (Fig.1). These two chimeric sequences were sent to the Genecust society (Luxembourg) to synthesize the corresponding CIMCs, but only pHIV-1/OM was successfully obtained.

**Figure 1:**
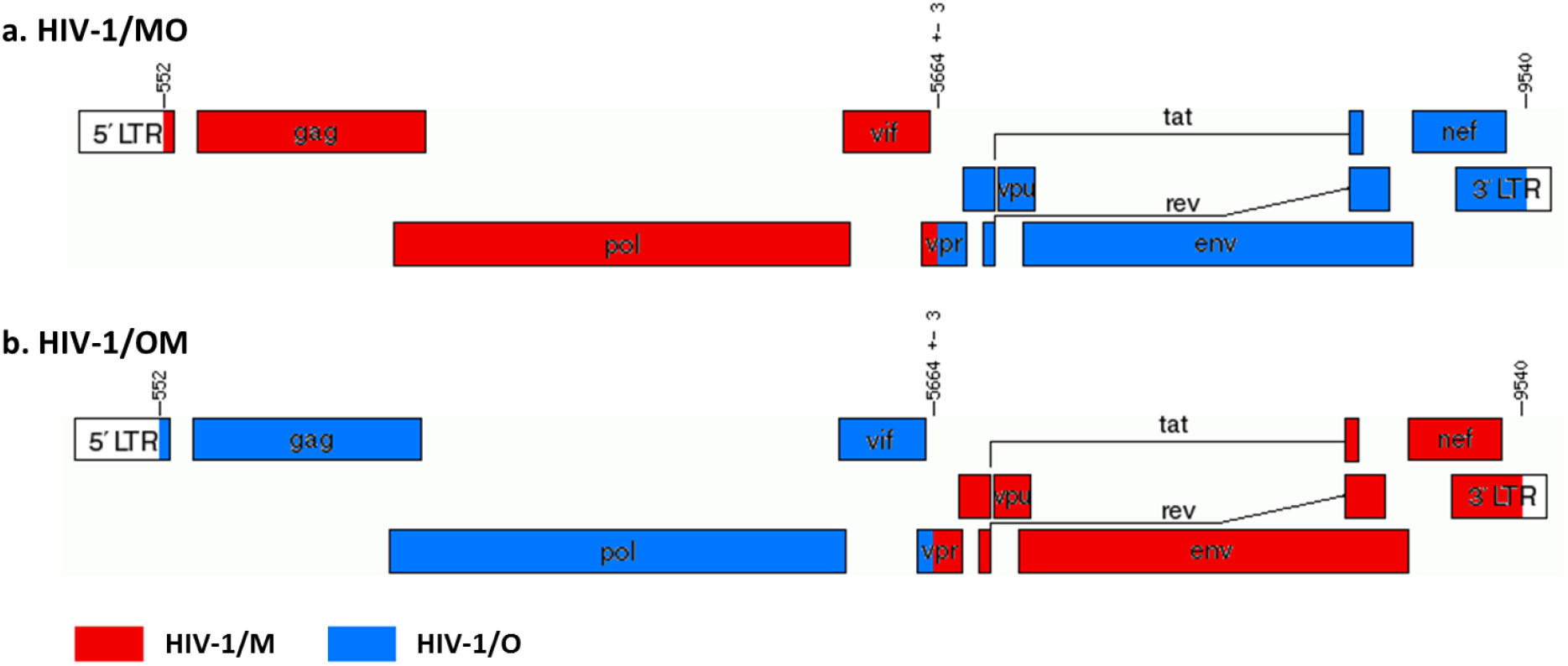
Representation of the HIV-1 intergroup M and O recombinant genomes

#### Cell lines

Human HEK-293T adherent cells were seeded at 37°C with 5% CO_2_ in DMEM + GlutaMAX™ (Gibco, Germany) containing 10% heat inactivated fetal calf serum (dFCS) (Eurobio Scientific, France) and gentamicin (50 μg/mL). Cells were maintained twice a week with trypsinization and one-third re-seeding. Jurkat.CD4-CCR5, obtained from the CFAR (NIBSC), is a CD4+ T non-adherent cell line with endogenous CXCR4 and stably transfected to express human CCR5. Cells were maintained in RPMI-1640 medium (Lonza, Swiss) supplemented with 10% dFCS, gentamicin (50 μg/mL) and geneticin G418 (500 μg/mL), at 37°C with 5% CO_2_, and were one-fifth re-seeded twice a week.

#### Production of parental and recombinant viruses from PIMC and CIMC

The parental and recombinant viruses were produced during a single experiment (Fig.2). For this, the two PIMCs (pRBF206 and p89.6) and the one CIMC (pHIV-1/OM) were rescued using a two-step reverse genetic system as follows. The day before transfection (D-3), 750 000 HEK293T cells were counted and seeded in a 6-well plate. The next day (D-2), HEK293T cells at 60-80% confluency were transfected with 1 μg of pHIV-1/M, pHIV-1/O, or pHIV-1/OM, using JetPrime™ (Polyplus Transfection, France), according to manufacturer’s instructions. At D0, a co-culture step was performed with 1.10^6^ Jurkat.CD4-CCR5 and at D2, supernatants were transferred to T25 flasks. Between D4 and D43, a fraction of each cell-free supernatant was removed, tested by quantifying p24 antigen (VIDAS^®^ HIV P24 II, BioMérieux, France), frozen and replaced by culture medium (RPMI + dFCS + gentamicin + geneticin), in order to remain at the same volume throughout the culture.

**Figure 2:**
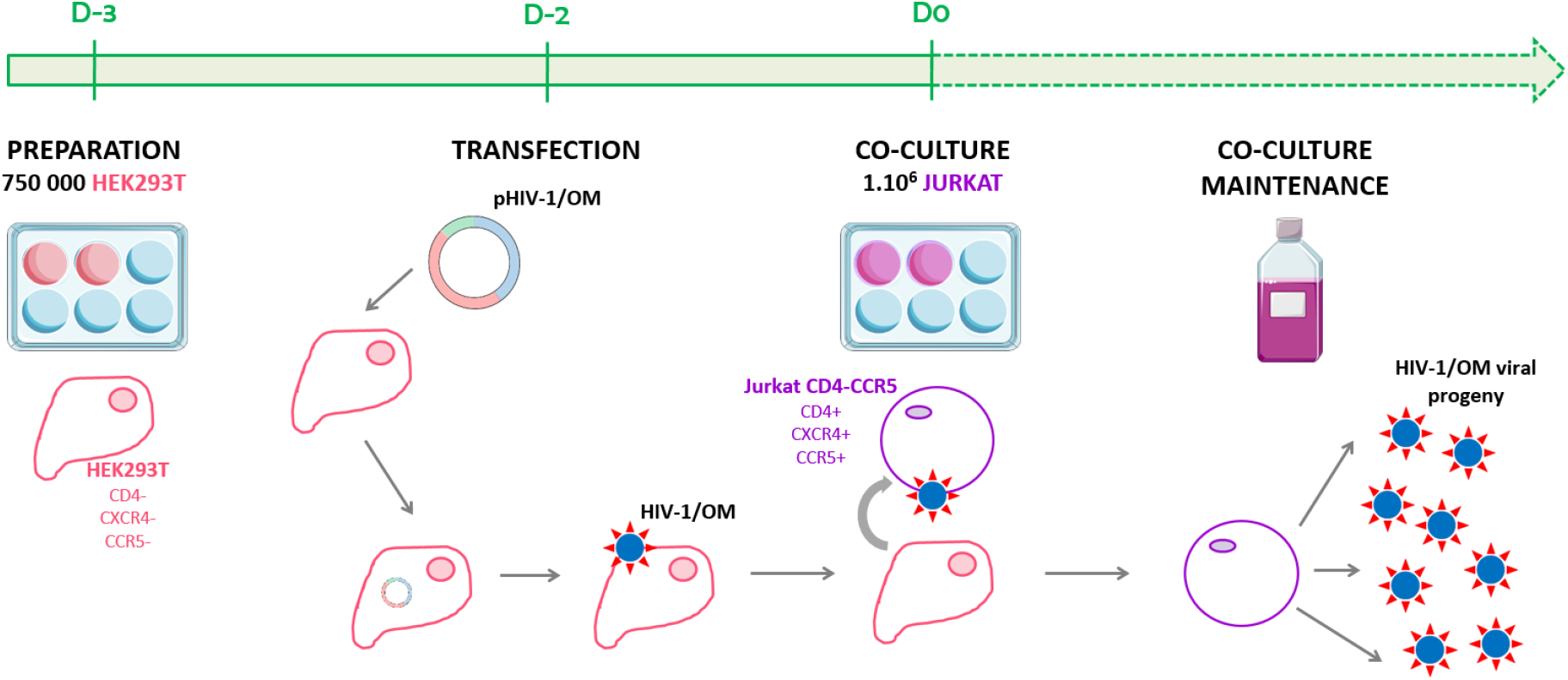
General principle of the production of HIV-1/OM recombinant viruses from a CIMC

The produced stock of HIV-1/M, HIV-1/O and HIV-1/OM supernatants was constituted by pooling if the concentration of p24 antigen was greater than 3 Log_10_pg/mL. The three viral pools were characterized by quantifying the p24 antigen (VIDAS^®^ HIV P24 II, BioMérieux), the viral load (Xpert^®^ HIV-1 Viral Load, Cepheid, USA) and the infectious virus titer by the 50% cell culture infectious dose (TCID50) Montefiori method, on TZM-bl cells. Finally, to eliminate any contamination of one virus by another, four nested RT-PCRs amplifying the Vif-Vpu region were performed as a quality control on each viral pool, as previously described [13]. The presence of a breakpoint in the *vpr* gene was investigated using homologous ([M-M] and [O-O]) and heterologous ([M-O] and [O-M]) combinations of HIV-1/M and HIV-1/O group-specific primers, followed by Sanger sequencing.

### Characterization of *in vitro* replicative potential

The replication kinetics of the parental and recombinant viruses was carried out in triplicate on the Jurkat.CD4-CCR5 line. For this, 5.10^6^ cells were infected with each virus (HIV-1/M, HIV-1/O or HIV-1/OM) at a multiplicity of infection (MOI) of 0.01, and in 5 mL of medium (RPMI + dFCS + Gentamicin + Geneticin). Between the first day (D1) and the last day (D30) of culture, 400 μL of supernatant were taken then frozen 6 days out of 7 and replaced by 400 μL of new culture medium, in order to maintain the same volume throughout the culture. The kinetics were measured on the basis of the RT activity (Lenti RT activity, Cavidi, Sweden) and the concentration of P24 antigen (INNOTEST^®^ HIV Antigen mAb, Fujirebio, Belgium) of each supernatant, for each triplicate of each virus, retrospectively. The mean values of the RT activity and of the concentration of P24 antigen of each triplicate were calculated (in Log_10_pg/mL) for each point of the kinetics and the peak was defined for each kinetic.

### Replication profile comparison

In order to highlight a possible difference between the HIV-1/OM recombinant virus and its HIV-1/M and HIV-1/O parental viruses, three parameters were considered: the time of reaching the peak (in days), the intensity of this peak (in Log_10_pg/mL) and the growth rate (in Log_10_/day). The growth rate was defined as the slope of the line connecting the point following the lowest point, reflecting the start of replication, and the highest point, corresponding to the replication peak.

## RESULTS

### CIMC pHIV-1/OM

The pattern of the HIV-1/OM recombinant virus involved RBF206 from the first bases of 5’LTR to the unique but conserved *AvrII* restriction site, in the *vpr* gene, and 89.6 from the *AvrII* site to the last bases of 3’LTR (Fig.1). The breakpoint in Vpr was localized at 5661-5667, relative to the reference form HIV-1/M HxB2 (GenBank accession number K03455). The CIMC pHIV-1/OM, synthesized by Genecust Society, corresponded to the HIV-1/OM full-length sequence inserted into a pET-28b(+) vector and was 15 358 bp in length (Fig.3).

**Figure 3:**
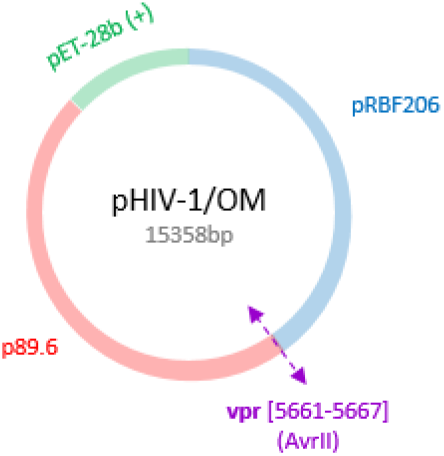
Profile of the CIMC pHIV-1/OM

### Production of parental and recombinant viruses from PIMC and CIMC

A pool of each virus (HIV-1/M, HIV-1/O and HIV-1/OM) was constituted and characterized. P24 antigen concentrations were between 4.5 and 5.9 Log_10_pg/mL, viral load values were between 8.8 and 9.3 Log_10_cp/mL and TCID50 values were between 91 376 and 267 184 IU/mL. (Table 1). These results showed very high viral production and reflected good infectivity for the three viruses considered.

**Table 1:** Characterization of parental and recombinant virus pools

|  | P24<br>antigen* | TCID50<sup>‡</sup> | VL<sup>°</sup> | Amplification of *vpr* gene |  |  |  | Pattern sequenced in *vpr* gene |  |  |  |
| --- | --- | --- | --- | --- | --- | --- | --- | --- | --- | --- | --- |
|  |  |  |  | [M-M] | [O-O] | [O-M] | [M-O] | [M-M] | [O-O] | [O-M] | [M-O] |
| HIV-1/M | 5.9 | 267184 | 9.3 | + | - | - | - | [M-M] |  |  |  |
| HIV-1/O | 4.5 | 226527 | 8.8 | - | + | - | - |  | [O-O] |  |  |
| HIV-1/OM | 4.5 | 91376 | 9.2 | - | - | + | - |  |  | [O-M] |  |

\*Log<sub>10</sub>(pg/mL)
‡TCID50/mL
°Viral load, Log<sub>10</sub>(cp/mL)

The RT-PCRs targeting the Vpr region confirmed the absence of contamination. Indeed, only the [M-M], [O-O] and [O-M] patterns were found for the parental (HIV-1/M and HIV-1/O) and recombinant (HIV-1/OM) viruses, respectively (Table 1). The [M-O] pattern, used as a negative control, was never amplified, as expected.

### Characterization of *in vitro* replicative potential

The triplicates of each virus showed good reproducibility, with only small differences calculated between the maximum and minimum values, both for the time to appearance of the peak and for viral production (Table 2). The recombinant virus showed a greater heterogeneity between triplicates than the parental viruses, with a difference of five days for the time to appearance of the peaks, and of 0.3Log_10_ and 0.8Log_10_ for the intensity of the peaks for RT activity and P24 antigen concentration, respectively. Finally, the kinetics of the two markers monitored, carried out independently in triplicate for each virus, proved to be reproducible and superimposable.

**Table 2:** Characteristics of the replication kinetics of each strain

|  | RT activity |  | P24 Ag |  |
| --- | --- | --- | --- | --- |
|  | Peak onset | Peak | Peak onset | Peak |
|  | time<br>(days) | intensity<br>*Log<sub>10</sub>(RT activity)* | time<br>(days) | intensity<br>*Log<sub>10</sub>(P24 Ag)* |
| HIV-1/M-1 | 15 | 5.1 | 12 | 6.7 |
| HIV-1/M-2 | 12 | 5.4 | 12 | 6.7 |
| HIV-1/M-3 | 15 | 5.1 | 11 | 6.7 |
| Mean value | 14 | 5.2 | 11.7 | 6.7 |
| Δmax-min | 3 | 0.3 | 1 | 0 |
| HIV-1/O-1 | 18 | 4.7 | 22 | 6.2 |
| HIV-1/O-2 | 17 | 4.9 | 25 | 6.3 |
| HIV-1/O-3 | 17 | 4.5 | 24 | 6.2 |
| Mean value | 17.7 | 4.7 | 23.7 | 6.2 |
| Δmax-min | 1 | 0.4 | 3 | 0.1 |
| HIV-1/OM-1 | 19 | 4.4 | 19 | 6.2 |
| HIV-1/OM-2 | 14 | 5.2 | 14 | 6.5 |
| HIV-1/OM-3 | 16 | 4.6 | 19 | 6.4 |
| Mean value | 16.3 | 4.7 | 17.3 | 6.4 |
| Δmax-min | 5 | 0.8 | 5 | 0.3 |

### Replication profile comparison

The mean kinetics of RT activity and P24 antigen concentration of the three viruses are shown in Figure 4a and 4b, respectively. The HIV-1/M parental virus grew faster and more intensively than the HIV-1/O parental virus, but also than the HIV-1/OM recombinant virus. Therefore, the supernatant samplings were stopped earlier for HIV-1/M than for the two other viruses (Fig.4). The replication kinetics of the recombinant virus appeared closer to its HIV-1/O parental virus than to its HIV-1/M parental virus (Fig.4). In order to quantitatively evaluate the differences between the three strains, we calculated the means of the data at the peaks of RT activity and of P24 antigen concentration of the triplicates and the growth rates as well as the differences for the three viruses (Table 3).

**Figure 4:**
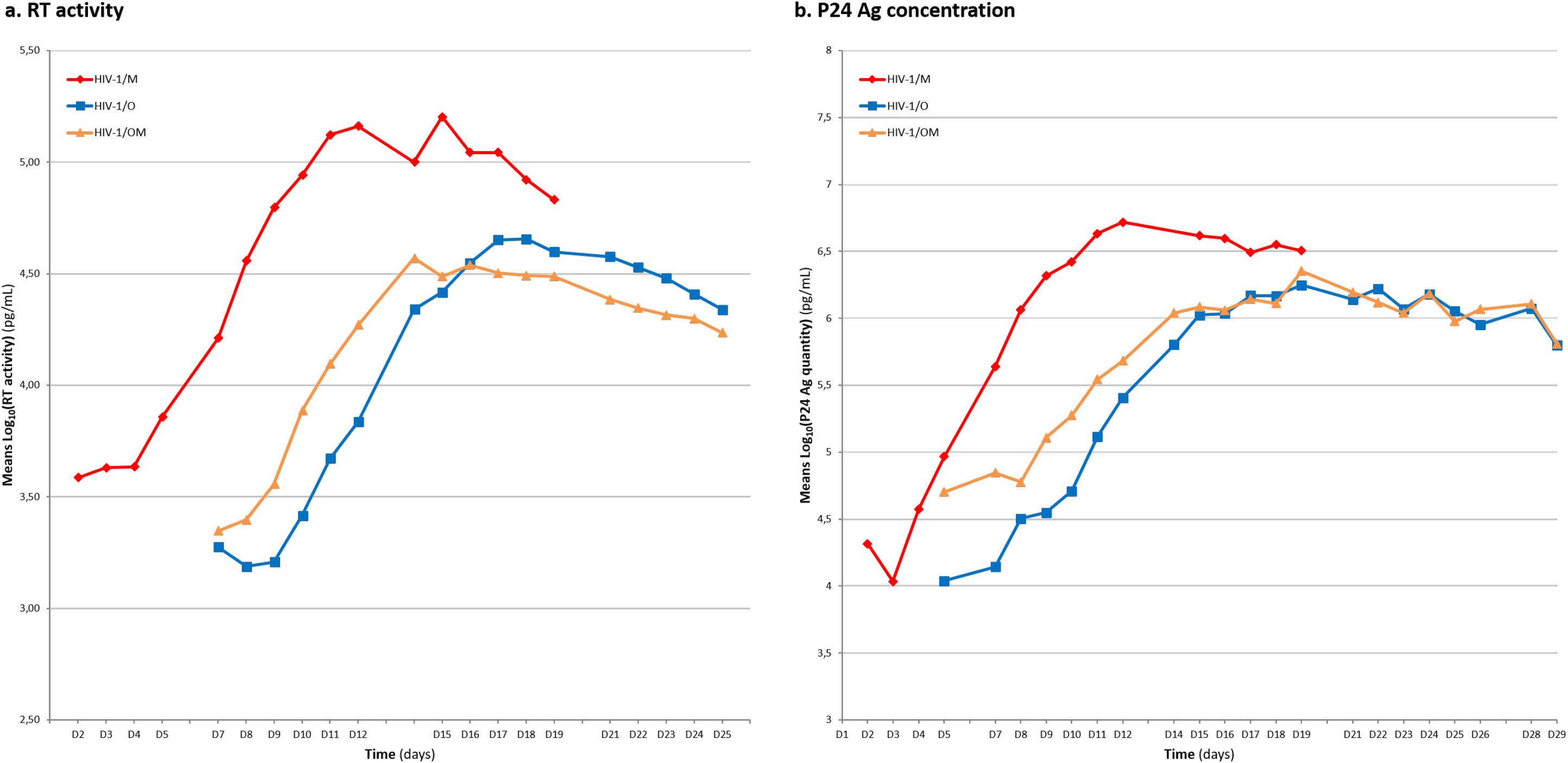
Mean kinetics of RT activity and P24 antigen concentration of parental and recombinant strains

**Table 3:**
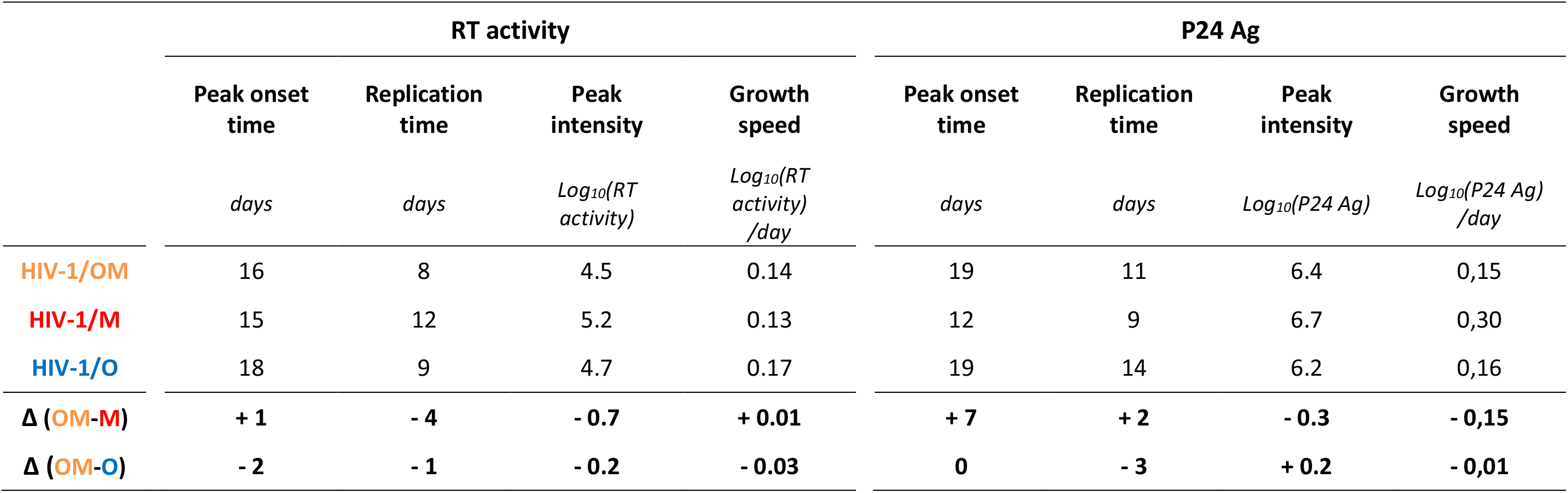
Mean values and differences calculated for the different parameters

This comparative analysis highlighted the hybrid behavior of the recombinant virus. Indeed, the peak of RT activity appeared earlier than for the parental virus HIV-1/O, which could be linked to the first steps of the viral replication cycle with an entry into the target cell favored by an envelope derived from HIV-1/M. On the other hand, concerning the intensity of RT activity and of P24 antigen production, the HIV-1/OM recombinant virus showed a behavior closer to that of the HIV-1/O parental virus, which could be similarly related to the recombination profile [O_gag/pol_-M_env_] we designed for our chimera (Table 3).

## DISCUSSION

A previous study suggested that an inter-group M and O recombinant form could be fitter than its parental forms [12]. Moreover, the abundant epidemic intragroup M recombinant forms, having emerged over time, are also in favor of this hypothesis of a better adaptation than the parental forms.

In this context, the objective of this study was to evaluate, for the first time, the *in vitro* replication of an inter-group M and O recombinant virus compared to its HIV-1/M and HIV-1/O parental viruses, to assess an increased potential for emergence in the human population.

Infectious molecular clones (IMCs) correspond to a full-length HIV genome sequence inserted into a vector and can produce virions, after transfection into susceptible cells. These newly formed virions all derive from a unique and known genetic sequence, as opposed to the high genetic diversity observed among quasispecies. The use of PIMCs to generate CIMCs, leading to the production of HIV-1/OM recombinant viruses by reverse genetics, allowed us to overcome any bias related to the use of heterologous strains, since the recombinant virus was compared to both the HIV-1/M and HIV-1/O parental strains, used for its generation.

Given that the HIV-1/M parental and HIV-1/OM recombinant viruses presented an X4 tropism in the V3 loop while the HIV-1/O parental virus presented an R5 tropism, the production of the viruses was carried out with the Jurkat CD4-CCR5 cell line. This cell line expresses the two co-receptors at its surface and can thus replicate viruses of different tropism. However, the proportions of expression of each co-receptor at the cell surface are unknown and an overexpression of one or the other could lead to a sensitivity of these cells to an infection depending on the tropism of the virus. In this context, we measured these expression percentages by flow cytometry. Among the Jurkat cells studied, 62.5% expressed both, whereas 35.3% and 1.5% expressed only the X4 and the R5 co-receptors, respectively. It is difficult to conclude as to the impact of the percentage of expression of the co-receptors on our comparative data with the HIV-1/O parental virus. However, our comparison between the HIV-1/OM recombinant virus and its HIV-1/M parental virus is validated since both of them share the same envelope and therefore the same tropism.

In order to eliminate any bias linked to the inter-manipulation variability of the cell culture, each kinetic of each virus was carried out in triplicate and the reproducibility of the values validated our comparisons. Overall, a few exceptionally outlying values were revealed, but the minimum and maximum values found in each triplicate remained close.

The choice of replication monitoring tools was decisive for our objective of comparing replicative potential. For P24 antigen production, the INNOTEST^®^ HIV Antigen mAb kit (Fujirebio) was used to avoid a possible misquantification linked to HIV-1/O antigenic diversity [21]. In addition, on each supernatant, RT activity was determined with the Cavidi method, which quantifies viral enzymatic activity without being impacted by viral diversity [22]. Moreover, the quantification of RT activity appears to be more representative of the replicative potential of the strain than the P24 antigen produced during replication, which corresponds to an accumulation of the protein over time.

The use of these two markers highlighted different behaviors depending on the virus and on the marker. Thus, the comparison of the kinetics of RT activity showed that the HIV-1/M parental virus replicated more quickly and more intensely than the HIV-1/OM recombinant and HIV-1/O parental viruses, but with an intermediate behavior of the recombinant virus. In fact, the results did not overlap completely either with those obtained with the HIV-1/M parental virus, or with the HIV-1/O parental virus. The P24 antigen production by the HIV-1/M parental virus was also greater than that of the other two viruses, with always an intermediate behavior of the HIV-1/OM recombinant virus.

Our results showed that the HIV-1/M parental virus had a significantly higher replicative potential than the HIV-1/O parental virus, as described *in vitro* by Arien *et al*. [19]. On the other hand, our results differed from the hypotheses put forward by Peeters *et al*. [12]. Indeed, after co-culture of PBMCs from a patient co-infected with an HIV-1/MO recombinant form and the HIV-1/M parental form from which it was derived, only the recombinant virus was detected. The team suggested a replicative superiority of the HIV-1/MO recombinant virus than its HIV-1/M parental virus [12], but the recombination profile described by Peeters *et al*. was [M_gag/pol_-O_env_], which could explain our discordant results. Thus, our results showed, whatever the marker, the clear replicative superiority of the HIV-1/M parental virus compared to the recombinant virus, even if the monitoring curves for RT activity and for P24 antigen production were not completely superimposable. However, the HIV-1/OM recombinant virus seems to have certain advantages over its HIV-1/O parental virus, but it was not possible, in our study, to demonstrate a systematic better replicative potential of the HIV-1/OM recombinant virus than its parental viruses.

This hybrid replicative behavior of our recombinant virus could be explained by the [O_gag/pol_-M_env_] recombination profile chosen. Indeed, its exponential growth phase began earlier than that of the HIV-1/O parental virus, suggesting a greater infectivity of the recombinant virus. The viral envelope is known to play a key role in the recognition of receptors and co-receptors, in the early phases of infection as well as in viral fitness [23, 24]. The envelope of the recombinant strain generated from the HIV-1/M parental virus could therefore be favorable to the recombinant virus. The predominance of group M envelopes among the HIV-1/MO recombinant forms described *in vivo* (7 profiles [O_gag/pol_-M_env_] versus 4 profiles [M_gag/pol_-O_env_] [15] supports this hypothesis. The production of the HIV-1/OM recombinant virus, on the other hand, appeared worse over time, possibly because the regions of the genome coding for the structural proteins and the replication enzymes came from the HIV-1/O parental virus. Indeed, the peak of RT activity of the recombinant strain was close to that of the HIV-1/O parental virus (lower by 0.2 Log_10_ pg/mL), as was the peak of the concentration of AgP24 (stronger by 0.2 Log_10_ pg/mL). The opposite pattern of recombination, [M_gag/pol_-O_env_], remains to be produced and tested, in order to confirm these hypotheses, unless this reverse virus is not viable, which is also one hypothesis.

Thus, these results should be linked to the diversity of inter-group M and O, which suggests that not all the recombinant forms are necessarily possible and that they do not necessarily all have the same profile. Due to the low availability of different PIMCs, in terms of genetic diversity, this work was carried out using a single representative of HIV-1/M, subtype B, associated with a single representative of HIV-1/O, subgroup T. To date, there is no HIV-1/MO recombinant form described *in vivo* that shows this association. Our previous study of all URF_MOs revealed that the genetic diversity of the mosaic fragments matched the molecular epidemiology in Cameroon, with a clear predominance of HIV-1/M CRF02_AG (48%) and of HIV-1/O sub-group H (84 %). Indeed, among these 19 URF_MOs, 8 (42%) combine HIV-1/M subtype CRF02_AG with HIV-1/O subgroup H and only 3 (16%) combine HIV-1/M subtype CRF02_AG with HIV-1/O subgroup T. This atypical combination of parental viruses might have had an impact on our results. Moreover, the RBF206 form did not seem to be the most representative of HIV-1/O, since its Vpu protein has the ability to counteract Tetherin, as described in HIV-1/M forms [25]. Its replicative potential, due to this particularity of Vpu, could therefore be greater than that of the other variants of the O group and explain the small difference in replication observed compared to the HIV-1/OM recombinant virus.

The HIV-1/OM recombinant virus produced and studied in this work may not be the most representative of the recombinant forms described *in vivo* but constitutes the first built and functional HIV-1/OM chimera. This work, based on our previous studies and our knowledge about HIV-1/MO recombination, offers many perspectives for studies, especially to evaluate the potential emergence of such recombinant HIV, taking into account genetic diversity in particular. Nevertheless, this HIV-1/OM recombinant virus has highlighted many hypotheses about the impact of recombination on replicative potential and has demonstrated once again the significant plasticity of HIV genomes and their infinite possibility of evolution.

## ACKNOWLEDGMENTS

We wish to thank the staff of the laboratory of Virology of Rouen University Hospital (especially Fanny Lermechain) and the members of the DYNAMICURE research team for the cloning experiments, especially Gaetan Riou, from Inserm UMR1234 - PANTHER Laboratory, for his help in flow cytometry experiments. We are grateful to Nikki Sabourin-Gibbs, Rouen University Hospital, for her help in editing the manuscript and to Denys Brand, from Tours University, for his rigorous proofreading.

## Supplemental Material (if necessary)

None

